# Phenotyping cowpea accessions at the seedling stage for drought tolerance using the pot method

**DOI:** 10.1101/2020.07.10.196915

**Authors:** Gabriel V. Nkomo, Moosa M. Sedibe, Maletsema A. Mofokeng

**Affiliations:** Department of Agriculture Central University of Technology Free State, South Africa; Agriculture Research Council Grain Crops, Potchefstroom, South Africa

**Keywords:** Accessions, cowpea, drought tolerance, phenotype, screen houses

## Abstract

One of the most important screening techniques used in cowpea selection for drought tolerance is screening at the seedling stage. The objective of this study was to phenotype 60 cowpea genotypes for seedling drought tolerance in screen houses (glasshouse and greenhouse). A triplicated 6 × 10 alpha lattice design with four blocks was used for the experiments. After planting, pots were watered to field capacity, thereafter watering was completely withheld for 4 weeks after planting (WAP), when plants were at the three-leaf stage. Principal component analysis revealed that of the 14 variables, the first four expressed more than 1 eigenvalue. Data showed that PC1, PC2, and PC3 contributed 39.3%, 15.2%, and 10% respectively, with 64.68% total variation. Bartlett’s test of sphericity was significant at *p*<0.05, while the Kaiser-Meyer-Olkin measure of sampling adequacy was 77. A PCA plot and biplot showed that the number of pods (NP), seeds per pod (SP), survival count (SC), pod weight (PWT), and stem wilting in week one (WWK1) had the most significant contributions to genetic variability to drought tolerance and to yield after stress imposition Based on the PCA, biplot, and cluster plot, the accessions IT 07-292-10, IT 07-274-2-9, IT90K-59, 835-911, RV 343, and IT 95K-2017-15 had the maximum variability in terms of number of pods, seeds per pod, survival count, pod weight and wilting in week one after drought imposition. Cowpea accessions 835-911, IT 07-292-10, RV 344, *Kangorongondo*, and IT 90K-59 were the major individuals that contributed mainly to domain information model (DIM) 1 and 2. The accessions that contributed the least were IT 89KD288, *Chibundi mavara*, and TVU12746. Thirty-six cowpea accessions from both screen houses were tolerant to drought, 15 were moderately tolerant, while 23 were susceptible. The findings of the study provided a useful tool for screening and determining drought-tolerant and susceptible accessions at the seedling stage. Thirty-six cowpea accessions from both screen houses were tolerant to drought as well as those that showed great variability can be used as parents in future cowpea breeding programmes.

## Introduction

Cowpea [*Vigna unguiculata* (L.) Walp.], Fabaceae, (2n = 2x = 22) is an important leguminous crop in developing countries, especially in sub-Saharan Africa, Asia, and Latin America, with a genome size of about 620 million base pairs (Boukar *et al*., 2018). The improvement of cowpea mainly dependent on breeding and selection from existing landraces according to the phenotypic variability, which is largely influenced by environmental conditions. According to the Food and Agriculture Organization of the United Nations (FAO), cowpea was grown on 1 million ha in Africa in 2014, with the bulk of production occurring in West Africa, particularly in Niger, Nigeria, Burkina Faso, Mali and Senegal (FAOSTAT, 2017). The global cowpea production was 5.59 million and the average yield 443.20 kg/ha (Gull *et al*., 2018). Africa leads in both cultivation area and production, accounting for about 95% of each. Niger and Nigeria are the leading producers of cowpea, together accounting for about 70% of the cowpea cultivated area and 67% of production worldwide. Most cowpea cultivars have relatively shorter growing period and maturation cycles of 60 to 80 days, which makes it suitable for drought-prone regions (Kyei-Boahen *et al*., 2017).

Drought is one of the most severe environmental stress, and it has a significant negative impact on crop yield. Gomes *et al*. (2019) recommend the use of water-efficient varieties in combination with good crop husbandry practices. Cowpea plants exposed to temperatures of 30 to 38 °C from eight days after emergence to maturity had very limited vegetative growth and reproductive potential (Singh *et al*., 2010). Hall e*t al*. (2003) observed that there is a great need to screen and breed for drought-tolerant and water use efficient varieties in Africa, as cowpea is grown mostly under rain-fed conditions, with frequent exposure to intermittent droughts. Gomes *et al*. (2019) recommend the use of well-adapted, early maturing cultivars in the smallholder farming sector to escape losses from late season droughts. Araujo *et al*. (2018) studied germination of cowpea cultivars under osmotic stress, seeds of three cowpea cultivars (BRS Tumucumaque, BRS Aracê, and BRS Guariba) grown at five osmotic potentials after three pre-treatments: pre-soaking in deionised water, pre-soaking in salicylic acid, and without pre-soaking. It was observed that salicylic acid promoted a reduction in abiotic stress, and BRS Guariba was more tolerant to water deficits and adjusted its cellular electrolyte leakage to increase its proline content under induced water stress.

In a wooden box experiment to screen cowpea recombinant inbred lines (RILs) for seedling drought tolerance, Alidu *et al*. (2018) used 200 inbred lines. It was observed that 12 RILS performed well for recovery, 13 RILS were susceptible to drought stress, and 11 RILS had higher relative water and chlorophyll contents. Ajayi *et al.* (2018) analysed 10 cowpea accessions under screen house conditions and observed significant differences among accessions for percentage plant recovery, stem regrowth, and stem greenness. For the evaluation of four Mozambican cowpea landraces for drought tolerance, Martins *et al*. (2014) determined that variability exists among the landraces in terms of growth under drought conditions, with *Timbawene moteado* having considerably higher leaf dry biomass, leaf and nodule protein content, and symbiotic nitrogen fixation compared to those of other landraces, as well as the lowest increase in proteolytic activity.

In a screen house experiment to select drought-tolerant cowpea seedlings, Ismai’la *et al*. (2015) evaluated 23 cowpea accessions at the seedling stage in the 2013 and 2014 growing seasons. They observed that plant height, number of leaves, and stem greenness were all affected by drought stress. It was found that five varieties, Kanannado, Danila, IT07K-297-13, IT03K-378-4, and Aloka local, were highly tolerant to drought. In addition, six varieties IT07K-322-40, IT07K-313-41, IT07K-291-92, IT06K-270, IT07K-244-1-1, and IT06K-275, were classified as highly susceptible to drought and the remaining 12 varieties were found to be neither tolerant nor susceptible to drought. Ismai’la *et al*. (2015) recommended the use of early maturing cowpea cultivars in order for farmers to escape the effects of a late season drought. Most cowpea plants exposed to moisture variation during the vegetative or reproductive stages perform poorly; hence, seedling-stage screening is ideal in this scenario. The objective of this study was to phenotype 60 cowpea genotypes for seedling drought tolerance in screen houses.

## Materials and Methods

### Plant material

A total of 60 cowpea accessions collected from three geographic origins were used in this study (Table 1). Out of these, 33 accessions were from the International Institute of Tropical Agriculture (IITA) in Nigeria, 19 accessions from the Agricultural Research Council – Grain Crops in South Africa, and eight accessions from smallholder farmers in Buhera District in Zimbabwe.

**Table 1:** List of cowpea accessions used in this study obtained from three geographic origin.

| **No** | **Name** | **Source** | **Origin** |
| --- | --- | --- | --- |
| **1** | Dr Saunders | ARC-GC | South Africa |
| **2** | IT96D-610 | IITA | Nigeria |
| **3** | RV 574 | ARC-GC | South Africa |
| **4** | RV 342 | ARC-GC | South Africa |
| **5** | Pan 311 | ARC-GC | South Africa |
| **6** | Bechuana white | ARC-GC | South Africa |
| **7** | Barapara jena | Buhera | Zimbabwe |
| **8** | TVU 9443 | IITA | Nigeria |
| **9** | 95K-589-2 | IITA | Nigeria |
| **10** | RV 344 | ARC-GC | South Africa |
| **11** | Agrinawa | ARC-GC | South Africa |
| **12** | IT 95K-207-15 | IITA | Nigeria |
| **13** | Orelo | IITA | Nigeria |
| **14** | TVU 9671 | IITA | Nigeria |
| **15** | Mutonono | Buhera | Zimbabwe |
| **16** | UAM-14-143-4-1 | IITA | Nigeria |
| **17** | 98K-503-1 | IITA | Nigeria |
| **18** | RV 503 | ARC-GC | South Africa |
| **19** | 86 D 1010 | IITA | Nigeria |
| **20** | TVU 9620 | IITA | Nigeria |
| **21** | RV 202 | ARC-GC | South Africa |
| **22** | RV 351 | ARC-GC | South Africa |
| **23** | Encore | ARC-GC | South Africa |
| **24** | TVU 14190 | IITA | Nigeria |
| **25** | IT 89KD-288 | IITA | Nigeria |
| **26** | RV 551 | ARC-GC | South Africa |
| **27** | IT 82E-18 | IITA | Nigeria |
| **28** | Barapara purple | Buhera | Zimbabwe |
| **29** | Kangorongondo | Buhera | Zimbabwe |
| **30** | 835-911 | IITA | Nigeria |
| **31** | ITOOK 76 | IITA | Nigeria |
| **32** | 98K-476-8 | IITA | Nigeria |
| **33** | Ziso dema | Buhera | Zimbabwe |
| **34** | Chibundi mavara | Buhera | Zimbabwe |
| **35** | 90K-284-2 | IITA | Nigeria |
| **36** | RV 221 | ARC-GC | South Africa |
| **37** | RV 343 | ARC-GC | South Africa |
| **38** | IT 98K-506-1 | IITA | Nigeria |
| **39** | Oleyin | IITA | Nigeria |
| **40** | IT 07-292-10 | IITA | Nigeria |
| **41** | IT 08K-150-27 | IITA | Nigeria |
| **42** | RV500 | ARC-GC | South Africa |
| **43** | IT 90K-277-2 | IITA | Nigeria |
| **44** | 98D-1399 | IITA | Nigeria |
| **45** | ITOOK 1263 | IITA | Nigeria |
| **46** | RV 563 | ARC-GC | South Africa |
| **47** | IT 18 | Buhera | Zimbabwe |
| **48** | RV 194 | ARC-GC | South Africa |
| **49** | 335-95 | IITA | Nigeria |
| **50** | TVU 12746 | IITA | Nigeria |
| **51** | IT 07-274-2-9 | IITA | Nigeria |
| **52** | 97K-499-35 | IITA | Nigeria |
| **53** | IT 07-318-33 | IITA | Nigeria |
| **54** | IT89-KD-288 | IITA | Nigeria |
| **55** | RV558 | ARC-GC | South Africa |
| **56** | IT 99K-573-2-1 | IITA | Nigeria |
| **57** | Mupengo dema | Buhera | Zimbabwe |
| **58** | CH47 | ARC-GC | South Africa |
| **59** | TVU 13004 | IITA | Nigeria |
| **60** | IT 90K-59 | IITA | Nigeria |

IITA: International Institute of Tropical Agriculture, ARC GC: Agriculture Research Council-Grain Crops

### Planting and data collection

Seeds of cowpea accessions were planted in 20 cm diameter pots filled with a mixture of topsoil and compost (3:1) in a greenhouse at the Agriculture Research Council – Grain Crops in Potchefstroom, South Africa, in January 2019 (environment 1). The experiment was repeated in a glasshouse in February 2019 (environment 2). An alpha lattice design with four blocks was used for both experiments. A total of 60 accessions were selected for drought tolerance at the seedling stage and were used in the experiments. A triplicated 10 × 6 alpha lattice design was used. The cowpea varieties were planted in 20 cm diameter pots in the screen houses. After planting, pots were watered to field capacity for establishment, thereafter watering was completely withheld for 4 weeks after planting (WAP), when plants were at the three-leaf stage. Thereafter, wilted plants of each variety were counted daily until all the plants of the susceptible lines appeared dead. Stress was measured by observing all dead plants in the susceptible group. Watering resumed at three weeks after stressing in both the greenhouse and glasshouse experiments until harvest. After the resumption of watering, numbers of recovered seedlings were rated for recovery. Based on the days to wilting and percentage recovery, the accessions were rated as either drought-tolerant or -susceptible.

### Data collection

#### i. Temperature conditions of the screen houses

The daily minimum and maximum temperatures of the screen houses were captured using temperature loggers. The loggers were placed in the screen houses and set to record the temperature at hourly intervals for the whole period of the experiment (Figure 1). The highest and lowest day temperatures recorded in the greenhouse (environment one) were 35.75 °C and 27.67 °C, respectively. The highest and lowest night temperatures recorded in the greenhouse were 26.87 °C and 19.99°C, respectively. The highest and lowest daytime temperatures recorded in the glasshouse (environment 2) were 36.4 °C and 19 °C respectively. The highest and lowest temperatures recorded in the glasshouse was 23.64 °C and 18.5 °C respectively.

**Figure 1:**
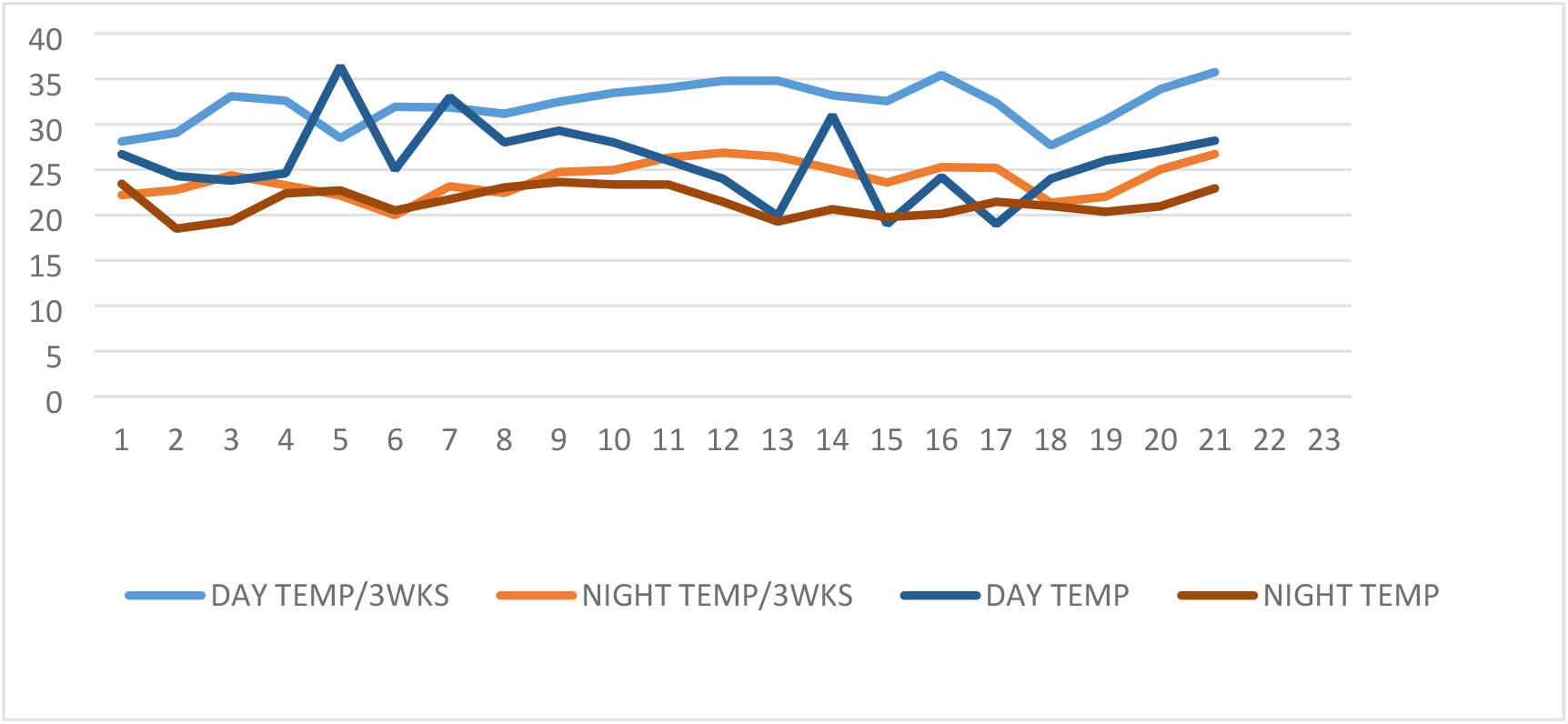
Graph showing day and night temperature ranges for 3 weeks.

#### ii. Agronomic traits

Drought tolerance was estimated using the wilting score (WS) as the degree of wilting severity, based on the 0–4 score scale as described by Singh *et al*. (2013). Recordings were done on seeds per plant, number of pods, number of seeds per pod, pod length, pod width and pod weight. Data were collected on number of days to seedling emergence, stem greenness, and wilting at 14, 21, and 30 days after planting (DAP), and rated on a scale of 0-–4 (Muchero *et al*., 2008).

Stem greenness

0 = leaves and stem completely yellow

1 = 75% of the leaves yellow, brown either from the base or tip of the stem

2 = 50% yellow or pale green, stem not turgid

3 = 25% yellow, 75% green, stem less turgid

4 = completely green, stem turgid

Wilting

0 = no sign of wilting

1 = 25% wilting

2 = moderate wilting, 50%

3 = yellow and brown leaves with 75% wilting

4 = completely wilted

After rewatering, data were collected on the survival count (SC): the number of surviving plants per genotype.

Recovery type

0 = no recovery

0.5 = recovery from the basal meristem

1 = recovery from the apical meristem

Recovery rate (RR)

The RR is computed as follows: (No. of dead plants/No. of emerged plants) × 100

## Data Analysis

A two-way analysis of variance (ANOVA) was used to determine significant differences in days to emergence (DTE), wilting scores, survival count, and yield-related traits. GenStat (version 19) software (www.genstat.kb.vsni.co.uk) was used for the statistical analysis of data. Statistical analysis was performed using IBM SPSS (version 20) (www.ibm.com/support/pages/spss-statistics-20-available-download) statistical computer package for principal component analysis, scree plot, and rotated component plot.

## Results

There were significant differences at the seedling stage of most genotypes tested on stem greenness in week one (SGWK1), stem greenness in week two (SGWK2), stem greenness in week three (SGWK3), wilting in week one (WWK1), wilting in week two (WWK2) and wilting in week three (WWK3) (Table S1). Most of yield-related traits evaluated were affected by water stress, with the exception of the environment on the number of pods (NP), environment on number of seeds per pod (NS), replicates on number of pods (NP), replicates on pod length (PL), and replicates on pod width (PWDTH) (Table S2).

### Principal component analysis

Principal component analysis (PCA) revealed that out of the 14 variables, all four components expressed more than 1 eigenvalue (Table 2). The proportion of variance among the four principal components (PCs) was 39.4% for PC 1, 15.2% for PC2, 10.1% for PC3, and 7.4% for PC4. The cumulative variance was 39.38% for PC1, 54.6% for PC2, 64.7% for PC3, and 72.1% for PC4. The first principal component (PC) was positively influenced by pod weight (PWT), with a value of 0.358, as well as by pod length (PL) (0.286), seeds per pod (SP) (0.263), seed weight (SWT) (0.255), and number of pods (NP) (0.181). PC2 was influenced by stem greenness at three weeks after planting (SGWK3), with a value measuring 0.332, and survival count (SC), with a value of 0.232. In PC3, stem greenness at week one (SGWK1) had the highest value (0.384), followed by stem greenness at week two (SGWK2) (0.295) and pod weight (PWT) (0.109). In PC4, days to emergence (DTE) had a positive influence (0.926), as did stem greenness at week two (SGWK2) (0.194).

**Table 2:** Eigen-values, proportions of variability and morphological traits that contributed to the first four PCs of cowpeas.

|  | PC1 | PC2 | PC3 | PC4 |
| --- | --- | --- | --- | --- |
| Eigen values | 5.510 | 2.130 | 1.410 | 1.040 |
| Proportion of variance (%) | 39.400 | 15.200 | 10.100 | 7.400 |
| Cumulative variance (%) | 39.380 | 54.600 | 64.700 | 72.100 |
| Day to emergence | 0.034 | 0 | -0.008 | 0.926 |
| SGWK1 | -0.050 | -0.101 | 0.384 | -0.169 |
| SGWK2 | -0.045 | 0.032 | 0.295 | 0.194 |
| SGWK3 | -0.123 | 0.332 | -0.004 | 0.088 |
| WWK1 | -0.070 | 0.130 | -0.339 | -0.039 |
| WWK2 | 0.069 | -0.117 | -0.216 | 0.063 |
| WWK3 | 0.111 | -0.277 | -0.052 | 0.004 |
| SC | 0.004 | 0.232 | -0.178 | 0.025 |
| Recovery Rate | 0.095 | -0.322 | 0.055 | 0.095 |
| NP | 0.181 | 0.063 | -0.036 | -0.047 |
| SP | 0.263 | -0.012 | -0.074 | 0.033 |
| PL | 0.286 | -0.048 | -0.074 | 0.035 |
| PWT | 0.358 | -0.283 | 0.109 | 0.022 |
| SWT | 0.255 | -0.034 | 0.027 | 0.012 |

SGK1 = stem greenness in week1; SGWK2 = stem greenness in week2; SGWK3 = stem greenness in week3; WWK1 = wilting in week1; WWK2 = wilting in week2; WWK3 = wilting in week3; SC= survival count; RR = recovery rate; NP = number of pods; SP = seeds per pod; PL = pod length; PWT = pod weight; SWT = seed weight.

A scree plot to show the relationship between eigenvalues and principal components was constructed to summarise the contribution of PCs (Figure S1). The plot showed that maximum variation was present in variable 1 with the highest eigenvalue of 5.8 followed by variable 2 (2.1), variable 3 (1.4), and variable 4 (1). Variable 14 had the lowest eigenvalue (0).

A further PCA with VARIMAX rotation was conducted to assess how 14 variables were clustered. The plot showed how closely related the 14 parameters were and these results are in tandem with the PCA plot (Figure S2). Three components were rotated based on the eigenvalues over 1 and the scree plot (Figure 1). Bartlett’s test of sphericity was significant at *p*<0.05 while the Kaiser-Meyer-Olkin measure of sampling of adequacy was 77, indicating sufficient items for each factor.

In the PCA plot, number of pods (NP), seeds per pod (SP), survival count (SC), pod weight (PWT) and wilting in week one (WWK1) had the most significant contributions to genetic variability in the drought tolerance cowpea accessions, as well as to yield after stress imposition (Figure 2).

**Figure 2:**
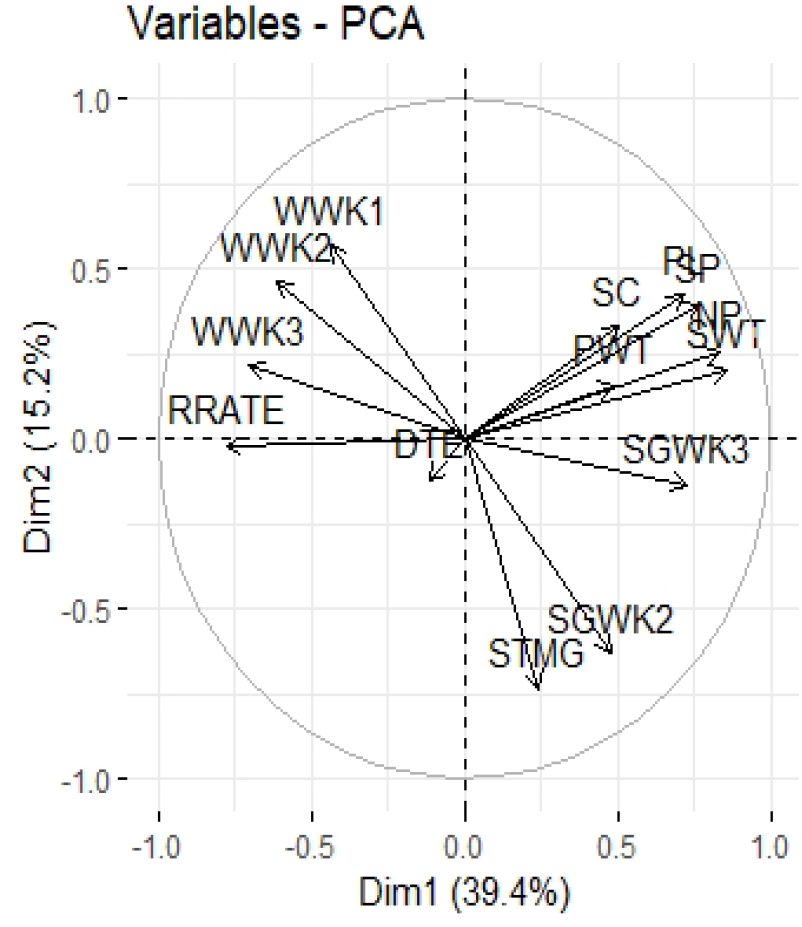
The contribution of various variables among 60 cowpea accessions screened for drought tolerance.

Cowpea accessions 835-911, IT 07-292-10, RV 344, *Kangorongondo*, and IT 90K-59 contributed the most to both domain information model (DIM) 1 and DIM 2 (Figure S3). The accessions IT 89KD288, *Chibundi mavara*, and TVU12746 contributed the least. The main contributors to DIM 3 were pod weight (PWT), stem greenness in week three (SGWK3), and recovery rate (RR) (Figure 3). The variables that contributed the least were seed weight (SWT), pod length (PL), and seeds per pod (SP).

**Figure 3:**
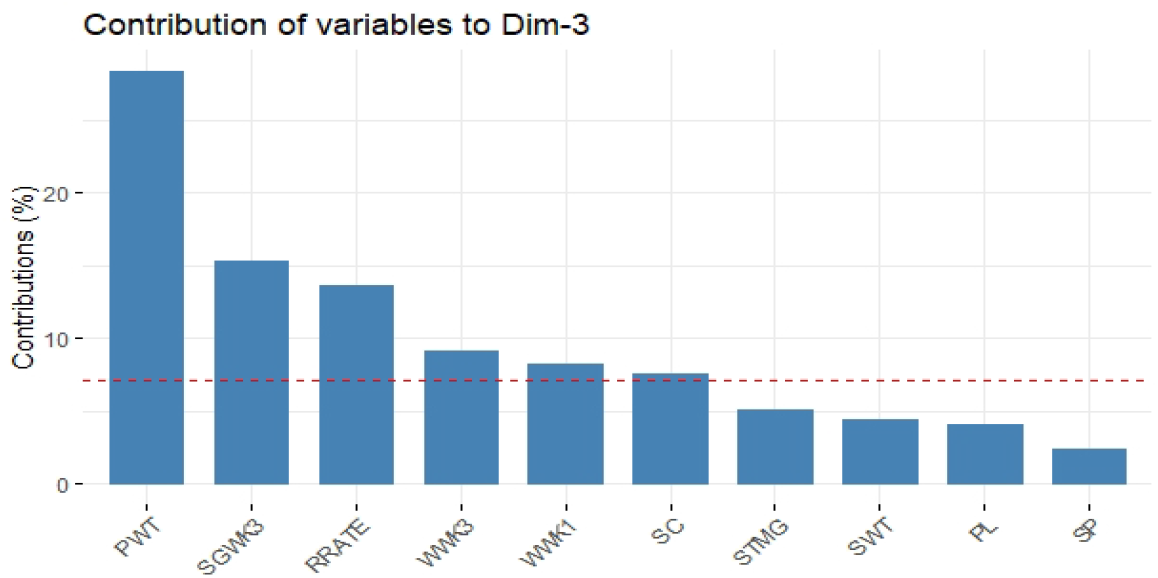
The contribution of various variables among 60 cowpea accessions screened for drought tolerance.

Figure 4 shows the relationships among traits in domain information model (DIM) 1 to DIM 5. Dim 1 was dominated by seed weight (SWT), number of pods (NP), and stem greenness in week three (SGWK3). DIM 2 was dominated by stem greenness in week one (SGWK1) and stem greenness at two weeks after drought imposition (SGWK2) and wilting in week one (WWK1). Pod weight (PWT) was the dominant trait in DIM 3, while in DIM 4 it was days to emergence (DTE).

**Figure 4:**
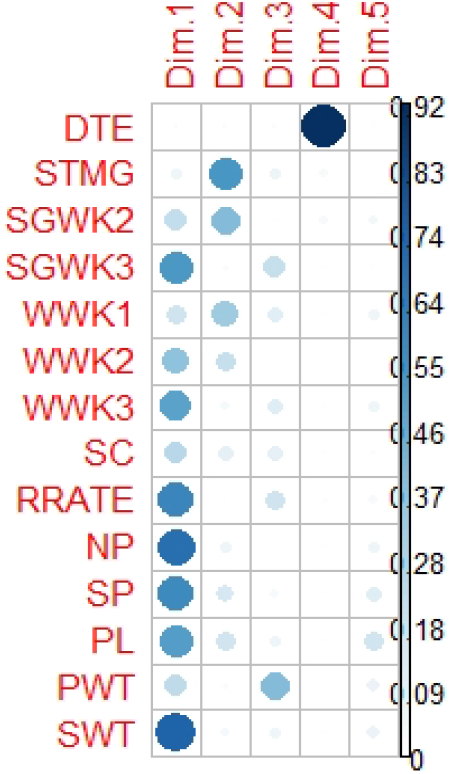
The contribution of various variables to Dim 1 to Dim 5.

The cluster plot analysis showed that the cowpea accessions can be grouped into three distinct clusters; red, blue, and green (Figure 5). Most accessions were grouped into the red and blue clusters. However, there was an overlap of accessions in the green and red clusters. As such, some accessions (TVU 13004, ITOOK 1263, IT89 KD 288, RV 588, Bechuana White, TVU 12746, IT07-318-33, and TVU 9671) managed to withstand water stress and flowered and produced pods when irrigation was resumed after three weeks of water stress.

**Figure 5:**
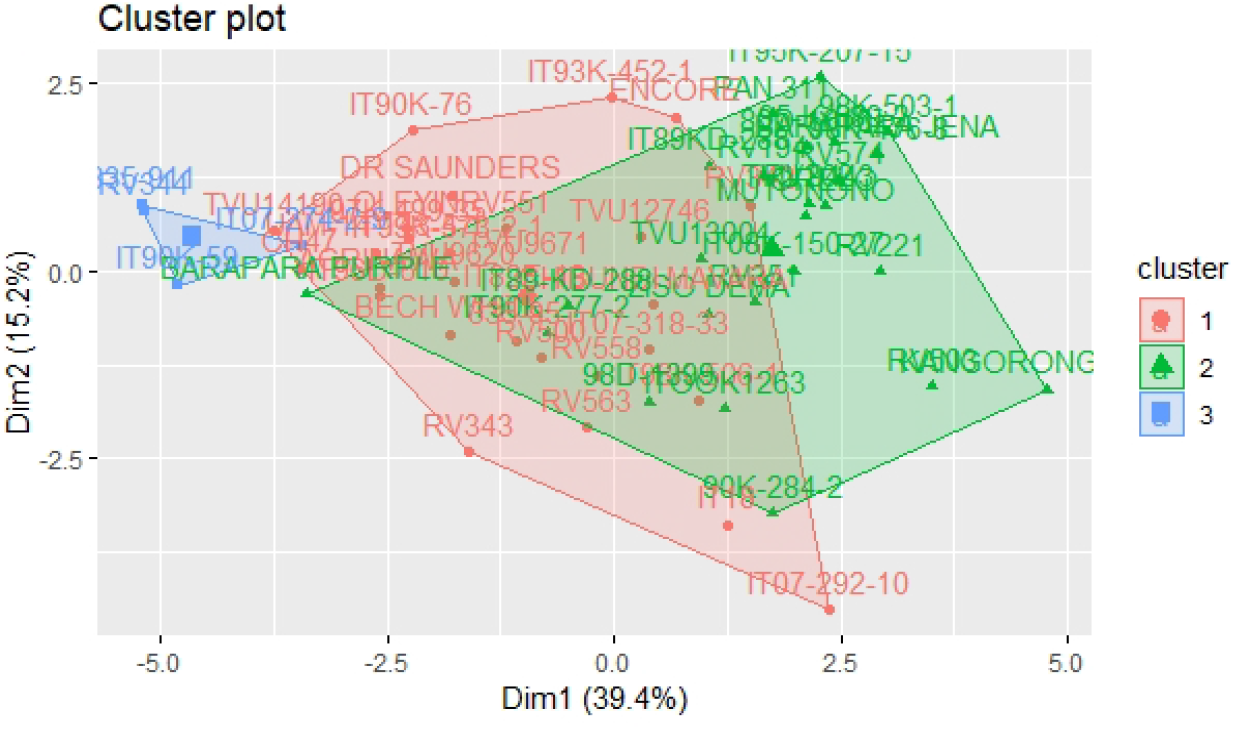
Cluster Plot showing the three groups of cowpea accessions grouped according to their levels of drought tolerance. Cluster 1 = Moderately tolerant, Cluster 2 = Susceptible and Cluster 3 = Tolerant.

The relationship of cowpea traits was studied using correlation coefficients. The correlation coefficient was weak and not statistically significant for SWT. However, the correlation coefficient was statistically significant in stem greenness in week two (SGWK2) compared to other traits. All of the significant correlation coefficients were positive and were mainly between days to emergence (DTE), stem greenness in week one (SGWK1), stem greenness in week three (SGWK3), survival count (SC), number of pods (NP), seeds per pod (SP), pod length (PL), pod weight (PWT), and seed weight (SWT) (Table S3). Pearson correlation analysis showed that the most significant relationships were observed from stem greenness in week two (SGWK2) up to seed weight (SWT). In addition, days to emergence (DTE) had mostly weak negative correlations with most of the measured attributes. Positive correlations were observed between seed weight (SWT) with stem greenness in week one (SGWK1), stem greenness in week two (SGWK2), stem greenness in week three (SGWK3), pod length (PL) and pod weight (PWT). Also there was positive correlation between pod weight (PWT) and stem greenness in week one (SGWK1), stem greenness in week two (SGWK2), stem greenness in week three (SGWK3), number of pods (NP), seeds per pod (SP) and pod length (PL). Pod length (PL) had positive correlations with stem greenness in week one (SGWK1), stem greenness in week three (SGWK3), number of pods (NP) and seeds per pod (SP).

The 60 cowpea accessions used varied in their response to drought imposition. Thirty-six cowpea accessions from both screen houses were tolerant to drought, 15 were moderately tolerant, while 23 were susceptible, based on the 14 traits measured (Table S4).

In the biplot, accessions IT 07-292-10, RV 343, and IT 95K-2017-15 had the maximum variability for the number of pods (NP), seeds per pod (SP), survival count (SC), pod weight (PW), and wilting in week one (WWK1) (Figure 6).

**Figure 6:**
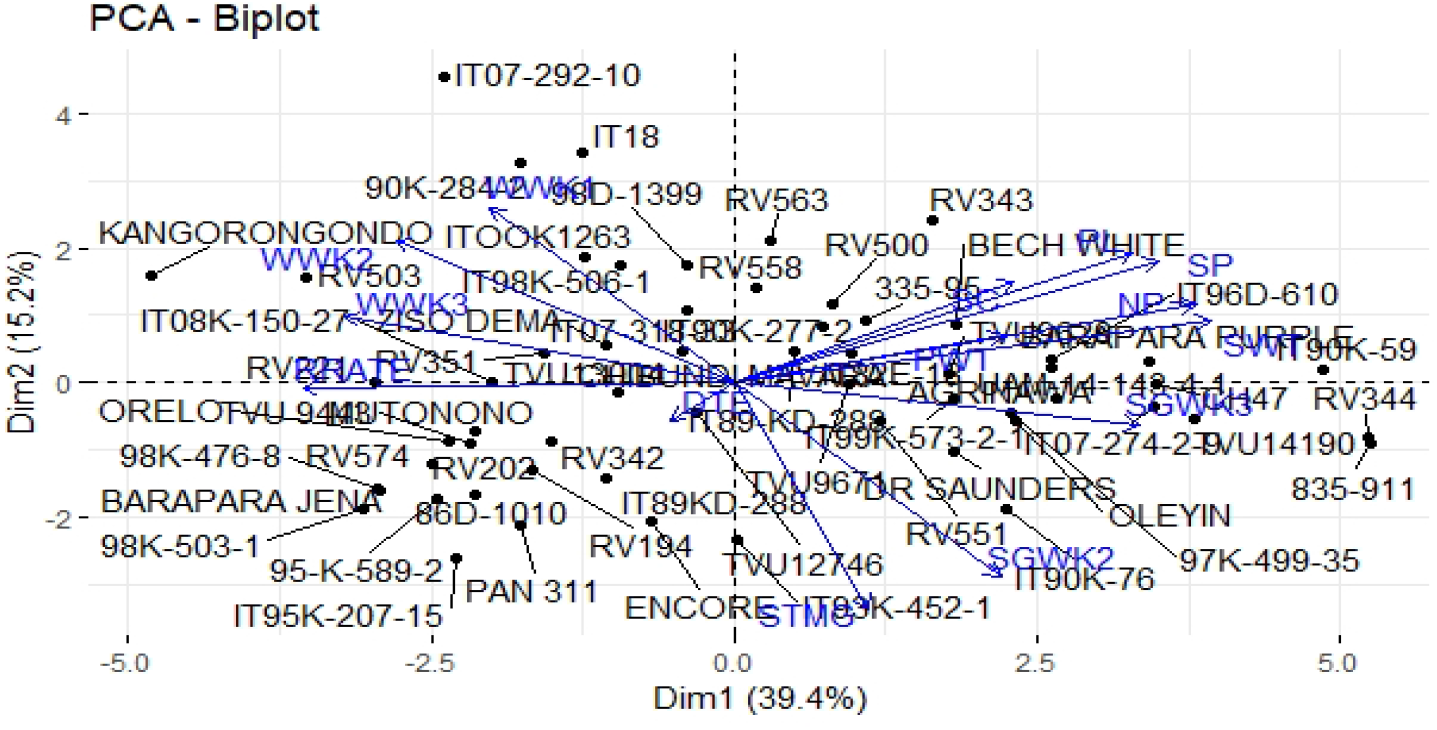
The contribution of various traits in the variability of 60 cowpea accessions to drought tolerance at the seedling stage.

The neighbour-joined cluster analysis generated by UPGMA divided the 60 cowpea accessions into two main clusters (Figure S4). From the results, it was observed that there were two major clusters and other sub clusters whose accessions were closely related genetically. The cluster analysis showed that the 60 accessions were grouped into two major clusters and other sub clusters with their respective distances. The phenotypic distance index based on morphological traits ranged from one (IT 89KD-288 from IITA) to 50 (TVU 13004 and IT96D-610 from IITA). The phenotypic distance index of other accessions in other subclusters was less than 20.

## Discussion

This study revealed that moisture is a very important component in plant growth and reproduction. According to Padi (2004), when moisture stress is imposed during the vegetative stage, it has the most effect on shoot and dry weight reduction in cowpeas. It is also during the vegetative stage that plants set up their architecture for reproduction. Alidu (2018) observed that moisture stress imposed after the pod-filling stage in determinate accessions has a limited reduction on the shoot and root biomass.

Most of the cowpea accessions showed differences in their response to drought imposition in their stem greenness from week one to week three after drought imposition. A similar variation was also observed when wilting was recorded from week 1 to week 3 after drought imposition. In both environments, the temperature had a significant effect on the performance of the accessions. In the greenhouse experiment, the average daytime and night-time temperatures were 34.24 °C and 23.98 °C, respectively. In the glasshouse experiment, the mean daytime and night-time temperatures were 26.06 °C and 21.42. According to DAFF (2011), the optimum temperature for growth and development of crops is around 30 °C. In this experiment, the 27 tolerant accessions were observed in the glasshouse. The relatively lower temperatures in the glasshouse might have accounted for the large number of drought tolerant cowpea accessions that were observed. This is agreement with Hamidou *et al*. (2007) who reported that water stress significantly increased canopy temperature of cowpea plants. Alidu and Pardi (2019), also confirmed that temperatures above 30 °C increases the intensity of stress levels in cowpeas. However even though Ndiso *et al*. (2016) observed that water stress significantly increased canopy temperature for cowpea varieties such as Khaki and Kaima-koko, it also reduced the canopy temperature of Macho and Nyekundu. Under higher temperatures in greenhouse, cowpea accessions such as IT 07-274-2-9, IT 90K-59, RV 344 and 835-911 perfomed better than any other accessions. These cowpea accessions ability to withstand drought and high temperatures is very important as they can be used as parents to study the inheritance of these specific responses in breeding.

Both the PCA plot and biplot highlight the importance of the distance of variables to PCs and their ultimate contributions to the drought tolerance of accessions, as well as to the yield after stress imposition. The PCA plot and biplot showed that the number of pods (NP), seeds per pod (SP), survival count (SC), pod weight (PWT), and wilting in week one (WWK1) had the most significant contributions to genetic variability in drought tolerance in cowpea accessions, as well as to the yield after stress imposition. In both the PCA plot and biplot, accessions placed far from each other were more diverse. Based on the PCA, biplot, and scatter plot, the accessions IT 07-292-10, RV 343, and IT 95K-2017-15 had the maximum variability for the number of pods (NP), seeds per pod (SP), survival count (SC), pod weight (PWT), and wilting in week one (WWK1), and could be used in future breeding programmes. These findings also conform with Al-Saady *et al*. (2018) who also used the PCA plot and biplot to reveal the large variation among 64 cowpea accessions in terms of seed length (cm) and width (cm), 100-seed weight (g), and seed colour.

In domain information analysis (DIM) 1 and DIM 2, seed weight (SWT), number of pods (NP), and stem greenness were the major determinants. Both groups had accessions 835-911, IT07-292-10, IT90-59, IT89KD288, *Chibundi mavara*, and TVU12746, which were tolerant to drought, while RV344 and *Kangorongondo* were susceptible to drought during the first week of drought imposition. Pungulani *et al*. (2012) also observed significant variations for 36 accessions that were characterised for canopy maintenance in a glasshouse. Only five accessions had apical re-growth, high relative water content, stem greenness and lower scores for both leaf wilting scales and leaf wilting index after the first week of stress at a soil moisture content of 2.9% while another five cowpea accessions showed high levels of drought susceptibility. Ravelombola *et al*. (2018) found large variations with three cowpea accessions that tolerant to wilting and drought on 30 cowpea accessions that were screened for drought-related traits at seedling stage in a greenhouse.

From this experiment, positive correlations were observed on stem greenness from week one up to week three as well as yield related traits such as seed weight (SWT), pod length (PL), pod weight (PWT), number of pods (NP) and seeds per pod (SP). Walle *et al*. (2018) observed significant and positive correlations among the number of pods per peduncle and number of seeds per pod, pod weight, seed length, seed thickness, seed weight, 100-seed weight, biomass, and harvest index at the genotypic and phenotypic levels. Diwaker *et al*. (2018) revealed that at the genotypic and phenotypic levels, a significant and positive correlation was shown by pod yield quintal per hectare with pod yield per plant and pod length. Ajayi *et al*. (2017) observed that the genotypic coefficient of variation (GCV) was lower than the phenotypic coefficient of variation (PCV) for all studied traits. They observed that both the GCV and PCV were reduced as drought stress went beyond 21 days among the wilting parameters and morphological traits, because of the influence of the environment on these traits. These findings are in tandem with results obtained from the study.

The main traits that accounted for variability from PC1 to PC5 in the screen houses were pod weight (PWT), pod length (PL), seeds per pod (SP), seed weight (SWT), number of pods (NP), stem greenness in week two (SGWK2), and days to emergence (DTE). This implies that accessions that emerged earlier and withstood the imposition of drought had higher chances of podding and producing seeds. Thus, it is imperative to consider these traits in further enhancing cowpea accessions’ tolerance to drought at the seedling stage. Ajayi *et al* (2018) recommends the drought susceptibility score, percentage of permanent wilting, stem greenness and regrowth, number of leaves, and stem girth as the most ideal traits for use in the study of drought tolerance in cowpea seedlings. However, Alidu, (2018) recommends a wide collection of cowpea lines in order to select the most tolerant genotypes for various growth stages as parents in a hybridisation programme.

On the cluster plot analysis, accessions in cluster 1 had higher values compared to all other clusters for all traits investigated in this study except for stem greenness in week one, two and three (SGWK1, 2 and 3) after drought imposition and wilting in week one (WWK1) after drought imposition. In both greenhouse and glasshouse experiments, this cluster had early maturing and high yielding accessions that can be used in future cowpea breeding programmes for drought tolerance at the seedling stage. The differences and similarities in accessions on some clusters as a result of their locations indicate the extent of accession exchange among farmers from different regions (Al-Saady *et al*., 2018).

## Conclusion

The findings of this study provided a useful tool for screening and determining drought-tolerant and -susceptible accessions at the seedling stage. The results of the investigation were also useful in selecting accessions especially for average seeds per pod (AVSPD), number of seeds (NS), pod length (PL), pod width (PWDTH), and pod weight (PWT) for further breeding programmes. Some accessions were able to perform well in both screen houses, under different temperature conditions. Based on the PCA, biplot, and cluster plot, the accessions IT 07-292-10, IT 07-274-2-9, IT90K-59, 835-911, RV 343, and IT 95K-2017-15 had the maximum variability for number of pods (NP), seeds per pod (SP), survival count (SC), pod weight (PWT), and wilting in week one (WWK1). Thirty-six cowpea accessions from both screen houses were tolerant to drought as well as those that showed great variability can be used as parents in future cowpea breeding programmes. This stability of accessions with minimal variation in any environment or location can serve as a genetic pool or germplasm collection for the breeding of drought-tolerant cowpea accessions.

## Acknowledgments

The project was supported by the Central University of Technology, Free State Research Grant Scheme as well as Agriculture Research Council Grain Crops, Potchefstroom.

